# RG203KR mutations in SARS-CoV-2 Nucleocapsid: Assessing the impact using Virus-like particle model system

**DOI:** 10.1101/2022.01.02.473343

**Authors:** Harsha Raheja, Soma Das, Anindita Banerjee, P Dikshaya, C Deepika, Debanjan Mukhopadhyay, Subbaraya G Ramachandra, Saumitra Das

## Abstract

The emergence and evolution of SARS-CoV-2 is characterized by the occurrence of diverse sets of mutations that affect virus characteristics, including transmissibility and antigenicity. Recent studies have focused mostly on Spike protein mutations; however, SARS-CoV-2 variants of interest (VoI) or concern (VoC) contain significant mutations in the nucleocapsid protein as well. To study the relevance of the mutations at the virion level, recombinant baculovirus expression system based VLPs were generated for the prototype Wuhan sequence along with Spike mutants like D614G, G1124V and the significant RG203KR mutation in Nucleocapsid. All the four structural proteins assembled in a particle wherein the morphology and size of the particle confirmed by TEM closely resembles the native virion. The VLP harbouring RG203KR mutations in nucleocapsid exhibited augmentation of humoral immune responses and enhanced neutralization by the immunized mice sera. Results demonstrate a non-infectious platform to quickly assess the implication of mutations in structural proteins of the emerging variant.

## Introduction

COVID-19 has been one of the leading causes of death globally since its emergence in December 2019. The coronavirus, SARS-CoV-2, has been identified as the causative agent, and it has a 30 kb single stranded genome encoding 4 structural and 16 non-structural proteins (1). During infection, SARS-CoV-2 virus enters the cells through the ACE2 receptor (present on the epithelial cells lining the respiratory tract), which is recognized specifically by the spike protein SARS-CoV-2 virus (2). Along with the spike protein (S), Envelope (E) and Membrane (M) glycoprotein together form the virion structure which surround the genomic RNA coated by the Nucleocapsid (N) protein. Studies have shown that these structural proteins elicit host immune response thereby generating specific antibodies against them (3, 4). Nucleocapsid has been shown to be highly immunogenic and a promising vaccine target in SARS-CoV infection as well (5, 6). New virus variants with mutations in these proteins are emerging continuously, with increased transmissibility and severity. It is of utmost importance to understand the molecular basis and effects of these mutations for an effective therapeutic and vaccine development. However, it is challenging to study them because of Biosafety level 3(BSL-3) requirement. We have designed Virus-like particle (VLP), which is composed of all the structural proteins that form non-infectious virus-like particles but generate immune responses similar to infectious virus particles enabling the study of mutation of all the structural proteins in a more physiologically relevant system. The VLP has been produced using Baculovirus mediated gene expression because of its advantages over Adenovirus and lentivirus systems (7, 8). The mutations, D614G and G1124V within spike and RG203KR within Nucleocapsid revealed plausible structural implications as depicted through previous studies (9, 10). D614G predominantly circulated worldwide and is presently incorporated into the backbone of all emerging strains (VoCs and VoIs). Clinical evidence has revealed to increase viral replication in the upper respiratory tract by augmenting infectivity and virion stability (11). G1124V is one of the major mutations on CD8 T cell epitopes in S protein, which might have significant implications in context to immunogenicity.

The R203K and G204R mutations in Nucleocapsid were first identified in the A2a lineage within China and subsequently have spread within other lineages across Western Europe, UK, and then to the US and other parts of the world through a number of VoCs and VoIs, viz., the Alpha, Gamma, Lambda and now in the most underscored VoC, the Omicron. This RG203KR mutation has been shown to enhance the infectivity, fitness and virulence (12-14). Recently, a different VLP approach has also been used to study the effect of Nucleocapsid mutations on transmissibility of the virus (15). However, the impact on immunogenicity remains to be studied. Here, we have incorporated these mutations to study their impact using VLP as a platform.

## Results and discussion

### Expression, purification and characterisation of SARS-CoV-2 VLP

We have expressed all the four structural proteins of SARS-CoV-2 in baculovirus expression system to form the VLP. These proteins were placed under separate promoters, cloned in the Baculovirus expression vector, BacPAK9 (Takara, USA) and transfected in Sf21 cells as described in the schematic (Fig. 1A), to yield recombinant baculovirus expressing SARS-CoV-2 structural proteins. Based on our earlier finding on the emerging mutations (9), we have generated three VLP constructs. First one contains sequences of the original Wuhan strain as prototype (WT-VLP). The second one harbours D614G and G1124V mutations in S protein (S mut-VLP). The third one harbours RG203KR mutation in N protein along with the previous S mutations (S+N mut-VLP). After Baculovirus titration, the expression of SARS-CoV-2 proteins through recombinant Baculovirus was confirmed 3-4 days post transduction of Sf21 cells by immunofluorescence (Fig 1B).

**Fig 1.**
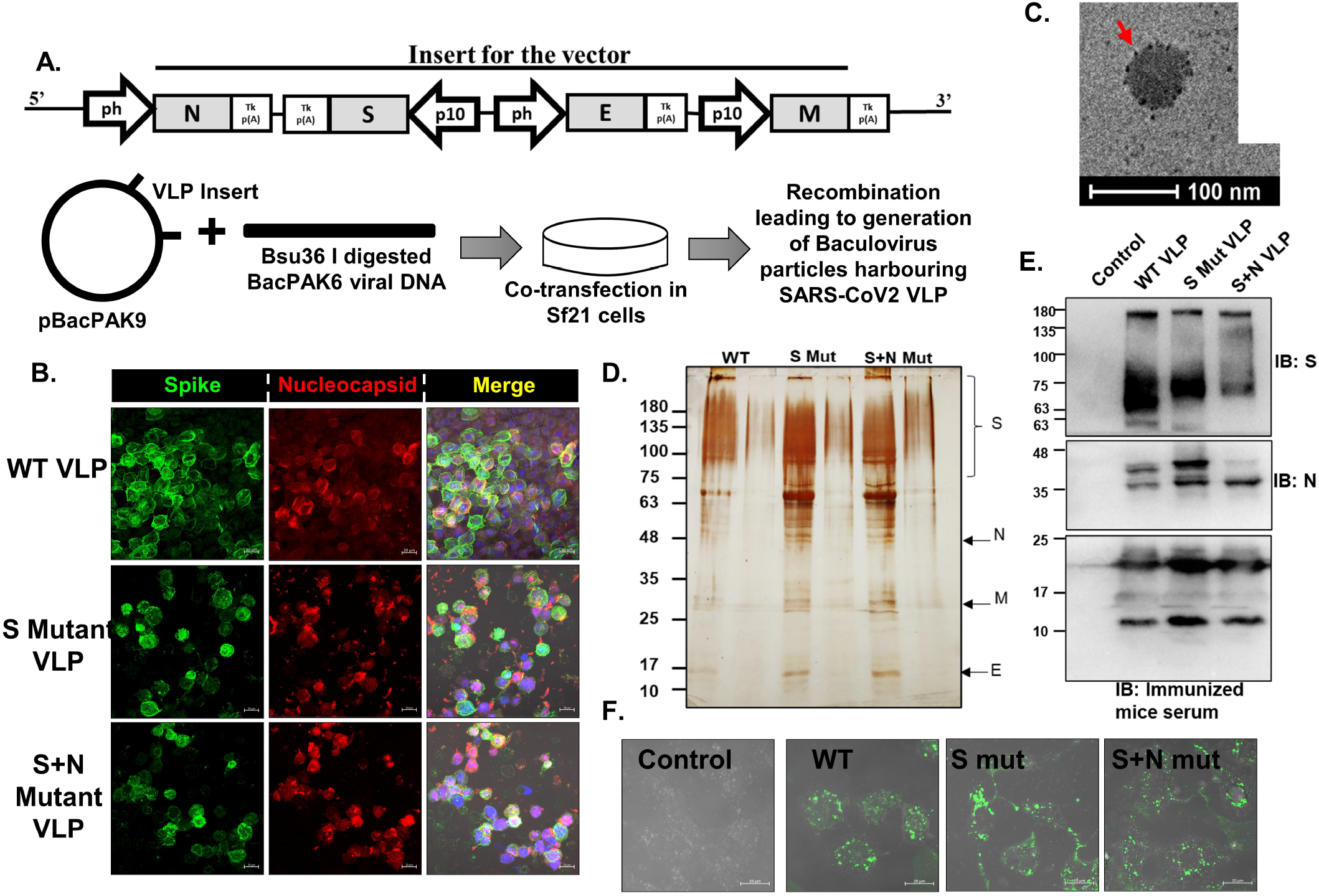
SARS-CoV-2 VLP purification and characterisation. **(A)** Schematic for the VLP expression construct and the baculovirus generation methodology. ph: Baculoviral-Polyhedrin promoter, p10: Baculoviral-p10 promoter, S: SARS-CoV-2 Spike protein, E: SARS-CoV-2 Envelope protein, M: SARS-CoV-2 Membrane protein, N: SARS-CoV-2 Nucleocapsid protein. **(B)** Baculovirus infected Sf21 cells were harvested after 96 h and processed for confocal staining using anti-S and anti-N specific primary antibodies and AF488 and AF633 labelled secondary antibodies. Nucleus was counterstained using DAPI. The bar represents 10um. **(C)**Transmission Electron Microscope images on purified VLP. The purified VLP was fixed and added to the copper grid, stained for S-protein using specific primary and immunogold labelled secondary antibody. Negative staining was done using uranyl oxalate. The arrow indicates immunogold labelled S protein. The purified VLPs were loaded onto SDS-10% polyacrylamide gel, followed by **(D)** silver staining (Adjacent to the VLP lane, a lower fraction of the gradient was loaded, confirming the VLP purity) and **(E)** western blotting to detect the presence of S-protein and N-protein using specific primary antibodies and HRP-tagged secondary antibodies. VLP injected mice sera was used as primary antibody followed by HRP-tagged anti-mouse antibody as secondary antibody. **(F)** Vero cells were incubated with AF488 labelled VLPs for 2h and processed for confocal imaging. The bar represents 20um.

VLPs were purified by overlaying the cell lysate 96 h post transduction, over 30-45%(w/w) sucrose gradient followed by ultracentrifugation at 28000 rpm for 3 h. VLP -containing band was used for characterisation. Transmission Electron Microscopy (TEM), involving negative staining and immunogold labelling for S protein, revealed the particle diameter in range of 30-100 nm (Fig. 1C). VLP purity was assessed by silver staining of the samples run on SDS-PAGE (Fig. 1D). 4 prominent bands around expected size of S (150-180 kDa), E (12 kDa), M (26 kDa) and N (48-49 kDa) proteins were observed, and further confirmed by Western Blotting using respective antibodies (Fig. 1E). Since anti-M and anti-E antibodies were not available commercially, sera obtained from mice injected with VLP was used to probe the blot.

To assess the physiological binding of VLPs to the ACE-2 receptor, Vero E6 cells were used. The VLP was fluorescence-labelled *in vitro* with Alexa Fluor 488 and its binding and internalisation was visualised by confocal microscopy (Fig. 1F). Concentration dependent increase of binding to Vero cells and absence of binding to ACE-2 deficient U937 cells confirmed the virus like, ACE-2 mediated, cell entry of VLPs.

### VLP induced immune response in mice

To assess their immunogenicity, the purified VLPs were injected in mice and sera collected as mentioned (Fig. 2A). We first checked the VLP tolerance by injecting a high dose (100 ug) in 6 weeks old Balb/c mice. We took 4 groups of mice, one for each VLP and one for vehicle control (PBS), with 6 mice in each group. The mice were monitored for 4 weeks for appearance of any toxic symptoms. All the mice survived with no effect on increase in body weight (Fig 2B). Additionally, administration of purified VLPs did not affect the histology of mice liver, kidney, heart and lungs as observed by histopathology (data not shown).

**Fig 2.**
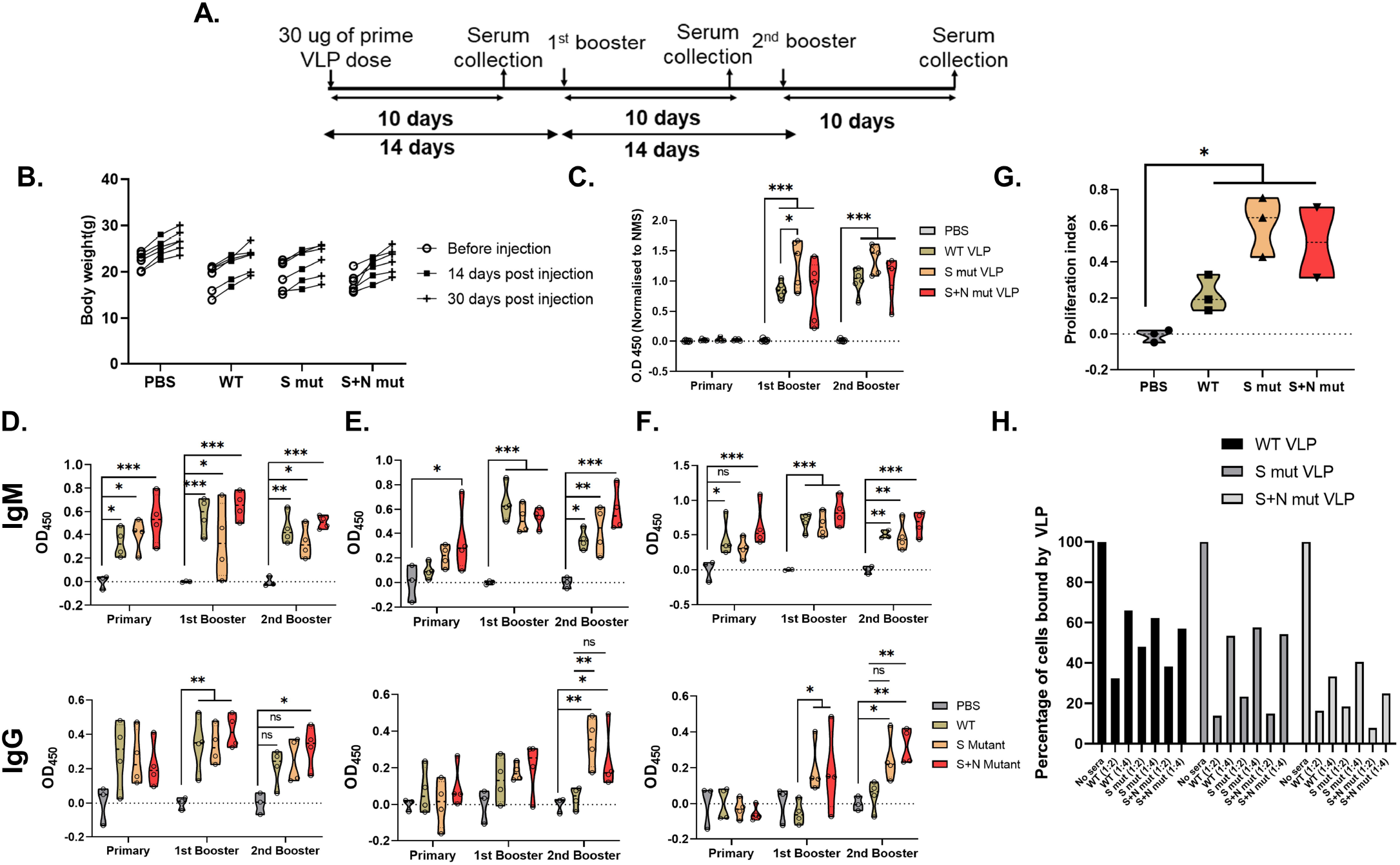
Immune response against SARS-CoV-2 VLP injection in mice. **(A)** Schematic of immunogenicity studies in mice. **(B)** Body weight of mice was measured at the indicated time points after VLP injection. ELISA was performed with murine sera collected after immunization with indicated VLPs at different time points. **(C)** WT-VLP, **(D)** Full length Spike protein **(E)** RBD of Spike protein **(F)** Nucleocapsid protein were used as antigen. Mice sera were added to the coated antigens and either HRP-tagged IgG+IgM (for panel C) or biotin labelled IgG/IgM antibodies (panels D, E and F) were used as secondary antibodies. Colour development by Streptavidin-HRP followed by addition of TMB substrate was quantified and plotted after normalisation as described previously. Two-way ANOVA was done for statistical analysis. p<0.05=*, p<0.01=**, p<0.001=*** **(G)** Splenocyte proliferation in response to peptides against Spike protein was quantified using MTT assay. MTT was added after 24 h of peptide stimulation and the colour development was quantified and plotted. Student’s t-test was done for statistical analysis. p<0.05=*, p<0.01=**, p<0.001=*** **(H)** Neutralisation of VLP binding to cells. Labelled VLP were incubated with indicated dilutions of sera prior to binding with Vero cells. VLP binding to cells after pre-incubation was analysed by flow cytometry and quantified.

Humoral response generated against the WT-VLP was quantified using ELISA and all the three VLP injected sera exhibited significant response as compared to vehicle control. Grossly, immunization with S mut-VLP elicited a higher response after the booster dose in contrast to WT or even S+N mut-VLP. (Fig. 2C). Humoral immunity in the form of IgM (first class of antibodies) and IgG (antibody produced after class switching) response was measured against total S protein (Fig. 2D), receptor binding domain (RBD) of Spike (Fig. 2E) and Nucleocapsid (Fig. 2F). Immunization with VLPs elicited a strong IgM and IgG response against total spike protein. Similar IgM response was observed against RBD and Nucleocapsid as well. Highest levels of IgM against all the three antigens were observed when immunized with the S+N mut-VLP, especially after the administration of booster doses. Interestingly, IgG response against RBD after second booster was much higher for S-mut VLP as compared to the WT-VLP. Interestingly, additional incorporation of N mutation reduced this response. For IgG response against Nucleocapsid after second booster, again there was heightened response in mutant VLP injected sera, and the incorporation of additional N mutation further increased the response. It appears that RG203KR mutation in nucleocapsid increases the IgG response against N, while reducing it against RBD, which indicates that the mutation in Nucleocapsid can potentially alter the viral structure, which could alter the antigenic sites on other proteins such as Spike. To further assess the T-cell response against the injected VLPs, proliferation of T-cells in response to *in vitro* stimulation with peptides against the S protein was measured using MTT assay. Significant difference in proliferation of T-cells isolated from VLP injected mice spleen as compared to vehicle control establishes the specific activation and proliferation of T cells by the injected VLP (Fig. 2G). Amongst the VLPs, as with the humoral response, mutant VLPs showed higher T-cell proliferation as compared to WT-VLP.

The highest titre sera obtained upon immunization with VLPs was further used to check the neutralisation of labelled VLP binding to Vero cells. We observed more than 50% VLP neutralisation in the presence of 1:2 dilution of sera from mice immunized with all the 3 VLPs (Fig. 2H). Notably, the efficiency of neutralisation of S mut and S+N mut-VLP was higher than the WT-VLP, pointing towards the accessibility of RBD in these VLPs for antibody neutralisation, as observed previously.

Taken together, we provide a comprehensive report of the impact of RG203KR mutation in nucleocapsid, on the immunogenicity and neutralisation efficiency using a model which can be easily manipulated and exploited to study the emerging SARS-CoV-2 mutations in a system closely resembling the virus while being non-infectious. It can be used to study the immune evasion capability of emerging viral variants and the efficacy of administered vaccines against those mutants. The mutations in structural proteins can easily be incorporated in the VLP which can be used to check neutralisation efficacy of sera from vaccinated individuals.

## Materials and Methods

### Cloning, transfection and generation of SARS-CoV-2 virus like particle (VLP)

The structural genes S, E, M and N with respective promoters were synthesized commercially (GenScript, USA) and cloned in pBacPAK9 with restriction sites BamHI and EcoRI. Vector DNA pBacPAK9 containing the target gene was transfected into *Spodoptera frugiperda* cells, along with Bsu36 I-digested BacPAK6 Viral DNA as described earlier (Takara Bio Inc., USA). Briefly, 1 × 10^6^ cells in 35-mm tissue culture incubated at 27°C for 1–2 hrs, washed with plain media and transfected with mixture containing DNA (100 ng/µl), Bsu36 I digested BacPAK6 viral DNA and Bacfectin and kept at 27°C for 5 hrs. 2% TC100 (Sigma, USA) media was added and kept at 27°C. ∼5 days after incubation, the medium, which contains viruses produced by the transfected cells, was collected and stored at 40C. The titres of the generated baculovirus were determined using BacPAK Baculovirus rapid titre kit (Clontech, USA).

### Immunofluorescence staining

For immunofluorescence staining of Sf21 cells infected with baculovirus expressing SARS-CoV-2 VLP, cells were seeded on coverslips in a 24-well plate for 14 h followed by infection with respective baculovirus. After the desired time of infection, cells were washed twice with 1X PBS and fixed using 4 % formaldehyde at room temperature for 20 min. After permeabilization by 0.1 % Triton X-100 for 2 min at room temperature, cells were incubated with 3 % BSA at 37 °C for 1 h followed by incubation with the indicated antibody for 2 h at 4 °C and then detected by Alexa-633-conjugated anti-mouse or Alexa-488 conjugated anti-rabbit secondary antibody for 30 min (Invitrogen). Images were taken using Zeiss microscope and image analysis was done using the Zeiss LSM or ZEN software tools.

### Transmission Electron microscopy (Immunogold labelling and negative staining)

The purified VLP was diluted in PBS, fixed with 4% paraformaldehyde and spotted onto 400 mesh carbon-coated copper grids for 10 min. It was then blocked using 1% BSA for 10 min, which was followed by incubation with primary antibody against SARS-CoV-2 S protein (Cat. No.-40592-R001) for 30 min. Thereafter, PBS wash was done 3-5 times and the grid was incubated with gold conjugated anti-rabbit secondary antibody for 15 mins. After 7-8 PBS washes, 1% glutaraldehyde was added onto the grid for 5 min to stabilise the immunostaining. Again, PBS wash was done 5 times and the samples negatively stained using 2% Uranyl oxalate. After thorough PBS washes, the grids were air dried and examined under transmission electron microscope at 80kV to visualise the immunogold labelled VLPs.

### Labelling of VLPs

SARS-CoV-2 LPs were labelled with Alexa fluor 488 carboxylic acid, succinimidyl ester (Cat. No.-A200000) using size exclusion chromatography columns. The labelled VLPs were used for binding with Vero cells in flow cytometry and imaging assays. For immunofluorescence imaging, Vero cells were seeded on coverslips in a 24 well plate. Labelled VLPs were added to the cells in DMEM media and incubated for 1-2 h at 37°C. Thereafter, coverslips were mounted on the slides and images taken in Zeiss710 confocal microscope and analyzed by Zen software tools.

### Isolation of VLP

The Baculovirus infected Sf21 cells were lysed with TEN buffer [10 mM Tris (pH 7.5), 1.0 mM EDTA, 1.0 M NaCl, 0.1% Triton X100, 1 mM PMSF]. For efficient lysis, the lysates underwent 2 freeze-thaw cycles in liquid Nitrogen followed by sonication at 3 sec on, 3 sec off for 2 minute cycle at 40 % efficiency setting. The lysates were centrifuged @3500 rpm for 30 minutes at 4°C. After centrifugation, the supernatant was collected and added on top of 30%-45% sucrose gradient and centrifuged @28000 rpm for 3 hours at 4°C in an ultracentrifuge using SW40 rotor. After centrifugation, the opaque band containing VLPs were collected and processed for characterisation.

### Western Blotting

Protein concentrations of the extracts were assayed by Bradford reagent (Bio-Rad) and equal amounts of cell extracts were separated by SDS-12 % PAGE and transferred onto a nitrocellulose membrane (Sigma). Samples were then analyzed by western blot using the desired antibodies, anti-SARS-CoV-2 S protein (Cat. No.-40591-T62), anti-SARS-CoV-2 N protein (Cat. No.-40143-MM05), Immunized mice sera followed by the respective secondary antibodies (horseradish peroxidase-conjugated anti-mouse or anti-rabbit IgG; Sigma). Antibody complexes were detected using the ImmobilonTM Western systems (Millipore).

### Animal immunization

Approval for animal experiments was taken from ‘Institutional Animal Ethics Committee’. Guidelines laid by the India National Law on animal care and use were followed for animal experiments. 24 female BALB/c mice, 6 weeks old, were grouped into four groups and immunization was given intra peritoneal (i.p.). SARS-CoV-2-LPs were conjugated with 2% alhydrogel as an adjuvant for immunization. In the first regimen, 30 μg of SARS-CoV-2-LPs was administered per mouse followed by two boosters with 15 μg SARS-CoV-2-LPs per mouse at an interval of 2 weeks between injection. In addition, mice group immunized with PBS served as a negative control. Pre-immune before the start of experiment and post-immune sera at each booster dose was isolated and stored at −70 °C. Mice were sacrificed and spleens removed at 10th day after final booster dose. Splenocytes were isolated as a mixed cell suspension using 70 μm cell strainer. ACK lysis buffer (155 mM NH_4_Cl, 10 mM KHCO_3_, and 0.1 mM EDTA) was used to deplete red blood cells from the cell suspension.

### Toxicity study in mice

24 male BALB/c mice (6 weeks old) were grouped into four groups and 100µg of SARS-CoV-2-LPs conjugated with 2% alhydrogel was administered by i.p. The weight and behaviour of the animals were monitored for 28 days. After 28 days, the animals were sacrificed and liver, lungs, heart and kidneys were extracted. To examine the toxicity effect four weeks post SARS-CoV-2-LPs administration in both control and injected groups, the histological analysis of 10% NBF fixed mice tissues were performed commercially.

### Measurement of Humoral Immune response after VLP immunization

ELISA was performed with murine sera collected after immunization with indicated VLPs at different time points. SARS-CoV-2 total spike proteins, receptor binding domains (RBD) and nucleocapsid (N) proteins were purchased from Sino Biologicals, China. ELISA was performed using Nunc MaxiSorp plates, Thermo Fisher Scientific, USA. Biotinylated goat anti mouse IgM, IgG, Streptavidin conjugated Horseradish peroxidase (Str-HRP), Bovine serum albumin (BSA) was purchased from Sigma Aldrich, USA. 3,3’,5,5’-Tetramethylbenzidine (TMB) solution was purchased from Applied Biological materials (ABM), Canada. All the rest of the chemicals were purchased from Sisco Research Laboratories (SRL), India and are of molecular biology grade.

ELISA was performed as described elsewhere (16). Briefly, ELISA plates were coated with 100 ng of proteins dissolved in PBS overnight. Next day, plates were washed with PBS with 0.05% Tween 20 (wash buffer) for three times and subsequently blocked with PBS with 2% BSA and 0.05% Tween 20 (blocking buffer) for 2 hrs. Thereafter 100 µl murine sera were added in 1 in 10000 dilution in blocking buffer for overnight. Following day, plates were washed thrice with wash buffer and 100 µl biotinylated goat anti mouse IgM (1 in 10000) or IgG (1 in 25000) was added to the wells for 2 more hrs. After 2 hrs, plates were again washed thrice with wash buffer and 100 µl streptavidin-HRP was added for 30 minutes. Finally, plates were washed five times with wash buffer and 100 µl TMB substrate was added. Following 10 minutes, 50 µl stop solution (2N HCl) was added and absorbance was recorded using a microplate reader (Spectramax M2e, Molecular Device, USA) at 450 nm. Statistical analyses were performed using Graph Pad Prism version 8.0. A Two-Way ANOVA followed by Tukey’s multiple comparison test was performed for determination of significance between groups and time points.

### Splenocyte proliferation assay

In a 96-well plate, 10^5 splenocytes were seeded and stimulated with peptides against S protein (From GenScript) (2μg/ml) in addition to CD28 for 24h. ConcavalinA (ConA) was used as a positive control. Proliferation was measured using 3-(4,5-dimethylthiazol-2-yl)-2,5-diphenyltetrazolium bromide (MTT). MTT was added to the splenocytes at a final concentration of 0.5mg/ml after 24h of peptide stimulation. After 3-4 h, media was removed, cells treated with 100µl DMSO and the absorbance measured at 560 nm. The proliferation index was calculated by using the following formula:

Proliferation upon stimulation= [O.D (Stimulated)-O.D (Unstimulated)] / O.D (Unstimulated)

Proliferation index= Proliferation upon peptide stimulation / Proliferation upon ConA stimulation

### Inhibition of binding of labelled VLPs to Vero cells by immunized mice sera

The labelled VLPs were incubated with 1: 2 and 1:4 dilutions of serum for 1 h at 37 °C. Vero cells (5×10^5^) were added to the mixture of SARS-CoV-2-VLPs and antibody in DMEM and 25 mM of HEPES buffer (100 µl) and incubated for 2 h at room temperature. Unbound complexes were removed by washes. Cell-bound fluorescence was analysed using an FACS Verse flow cytometer (Becton Dickinson) using BDFACSuite software to calculate the cell population bound by VLPs and percent binding was determined from the equation:

% Binding of VLP to cells = [percentage of VLP bound cells in experimental sample−negative control (only cells)] / [percentage of VLP bound cells in positive control (no sera) −negative control (only cells)] ×100.

## Acknowledgements

This work was supported by the Department of Biotechnology (DBT), (BT/PR40824/COV/140/3/2020); DBT-IISc Partnership Program; Department of Science and Technology, (DST-FIST).

S.D acknowledges J.C. Bose Fellowship from DST. H.R is supported by Council of Scientific and Industrial Research, India (SPMF). So.D is supported by WoSA, DST. A.B is supported by NPDF, DST.

## Notes

### Competing Interest Statement

The authors have declared no competing interest.

